# Divergent lineages in a young species: the case of Datilillo (*Yucca valida*), a broadly distributed plant from the Baja California Peninsula

**DOI:** 10.1101/2023.05.22.541794

**Authors:** Alberto Aleman, Maria Clara Arteaga, Jaime Gasca-Pineda, Rafael Bello-Bedoy

## Abstract

**Premise:** Globally, barriers triggered by climatic changes have caused habitat fragmentation and population allopatric divergence. Across North America, oscillations during the Quaternary have played important roles in the distribution of wildlife. Notably, diverse plant species from the Baja California Peninsula, in western North America, exhibit strong genetic structure and highly concordant divergent lineages across their ranges, as they were isolated during the Pleistocene glacial-interglacial cycles and thus accumulated genetic differentiation in their genomes. A representative plant genus of this Peninsula is *Yucca*, with *Yucca valida* having the widest range. Whereas *Y. valida* is a dominant species, there is an extensive distribution discontinuity between 26° N and 27° N, where no individuals have been identified, suggesting restricted gene flow. Moreover, the historical distribution models indicate the absence of an area with suitable conditions for the species during the Last Interglacial, making it an interesting model for studying genetic divergence.

**Methods:** We examined the phylogeography of *Y. valida* throughout its range to identify the number of genetic lineages, quantify their genetic differentiation, reconstruct their demographic history and estimate the species’ age.

**Key results:** We assembled 4,411 SNPs from 147 plants, identifying three allopatric lineages. Our analyses support that genetic drift is the driver of genetic differentiation among these lineages. We estimated an age under one million years for the common ancestor of *Y. valida* and its sister species.

**Conclusions:** Habitat fragmentation caused by climatic changes, low dispersal, and an extensive geographical range gap acted as cumulative mechanisms leading to allopatric divergence in *Y. valida*.

## INTRODUCTION

Genetic divergence, the formation of two or more independent lineages from an ancestral one, can be understood by two broad non-mutually exclusive frameworks: allopatric divergence, in which low dispersal, demographic fluctuations, or population extrinsic barriers lead to genetic differentiation (Mayr et al., 1963; Barton, 2008; Scheiner and Mindell, 2020) and ecological divergence, in which selection drives differentiation by cumulatively restricting the exchange of certain loci (Darwin, 1859; Tigano and Friesen, 2016; Stankowski and Ravinet, 2021). Identifying how genetic differentiation is accumulated between and within species is fundamental for understanding their evolutionary histories (Han et al., 2017; Irwin et al., 2018; Shang et al., 2022) and permits linking these histories to geographical regions’ geologic and climatic histories (Avise, 2000).

Globally, barriers triggered by climatic changes have caused habitat fragmentation and population allopatric divergence (Kumar and Kumar, 2018). For example, climatic oscillations during the Quaternary have played important roles in the distribution of wildlife across North America (Shafer et al., 2010), causing genetic differentiation in numerous taxa (Hewitt, 2000; e.g., de Oliveira et al., 2021). Likewise, since these climatic oscillations have been cyclic, divergence followed by a secondary contact due to habitat relinkages has been exposed as a common process in the Northern Hemisphere (e.g., Harrington et al., 2018; Rödin-Mörch et al., 2019).

Broadly distributed plant species are valuable models for studying genetic divergence (Petit and Hampe, 2006). Under wide geographic ranges, plant populations are expected to be particularly subject to demographic changes related to genetic drift (e.g., habitat fragmentation) and gene flow (e.g., changes in their dispersers dynamics) (Sork et al., 2016). Notably, diverse plant species from the Baja California Peninsula (BCP), located in western North America, exhibit strong genetic structure and highly concordant differentiated lineages across their ranges. Populations of cacti, ocotillos, palms, trees, and agaves from the BCP were isolated during the Pleistocene glacial-interglacial cycles and accumulated genetic differences in their genomes (Clark-Tapia and Molina-Freaner, 2003; Fehlberg and Ranker, 2009; Rebernig et al., 2010; Lira-Noriega et al., 2015; Gutiérrez-Flores et al., 2016; Klimova et al., 2017; Martínez-Noguez et al., 2020; Pérez-Alquicira et al., 2023). This is because during the glacial periods, the deserts in the Northern Hemisphere increased their distribution as the climate was drier, while the interglacial periods were highly humid, producing a significant fragmentation of these ecosystems (Qiu et al., 2011; Meng et al., 2015). Collectively, these patterns of genetic differentiation throughout the BCP indicate that divergence in this region has been strongly driven by allopatry (Dolby et al., 2015, 2022).

A representative plant genus of the BCP is *Yucca* (Asparagaceae), a monocotyledonous, long-lived, semi-evergreen tree with woody stems, lanceolate succulent green leaves, and white paniculate inflorescences. It is characterized by a long life cycle and slow population growth and recruitment rates (Turner et al., 2005). Its pollination depends on an obligate mutualism with moths of the genera *Tegeticula* and *Parategeticula*, which disperse pollen over short distances (Pellmyr, 2003), while seed dispersal has been related to extinct megafauna and is currently mediated by rodents, birds, and ruminants (Lenz, 2001). Three *Yucca* species are distributed across the BCP: *Yucca schidigera*, *Yucca capensis,* and *Yucca valida*, the latter two being endemic and sister taxa that are pollinated by a single moth species, *Tegeticula baja* (Pellmyr et al., 2007, 2008). Among the *Yuccas* of the BCP, *Y. valida* has the widest range, from ∼29.8° N to ∼25.6° N, across the Central and Vizcaino deserts, and the Magdalena Plains (Turner et al., 2005; Thiede, 2020). Whereas populations of *Y. valida* are continuous across their northern range, there is an extensive gap between 26° N and 27° N (our personal observation), where no individuals have been identified, suggesting restricted gene flow. Furthermore, while *Y. valida* is a widespread taxon, the historical distribution models indicate the absence of an area with suitable conditions for the species during the Last Interglacial (Arteaga et al., 2020). As a result, this species is an interesting model for studying genetic divergence.

We examined the phylogeography of *Y. valida* throughout its range. We hypothesized that its populations have experienced allopatric divergence, and since this species has a short-distance dispersal that limits genetic interchange between distant populations (Alamo-Herrera et al., 2022), we expected to find differentiated genetic lineages across its distribution. We aimed to i) identify the number of genetic lineages of *Y. valida* throughout its range, ii) quantify the levels of genetic differentiation between these lineages, iii) reconstruct their demographic history and iv) estimate the species’ age. This information will contribute to the understanding of the processes driving genetic divergence in the BCP and Western North America.

## MATERIALS AND METHODS

### Sampling, DNA extraction and sequencing

Fresh leaf tissue samples of 120 plants were collected from 20 locations across the species range (Figure 1; Appendix S1, Table S1 – see the Supplementary Data with this article). All samples were dried in desiccant silicate gel. We used 100 mg of each dried sample for DNA extraction following the Qiagen DNeasy Plant Mini Kit protocol (Qiagen, Hilden, Germany). The extractions’ quality and concentration were evaluated using GelRed 1.5% agarose gels and a NanoDrop spectrophotometer (Thermo Fisher Scientific, Waltham, Massachusetts, USA).

**Figure 1.**
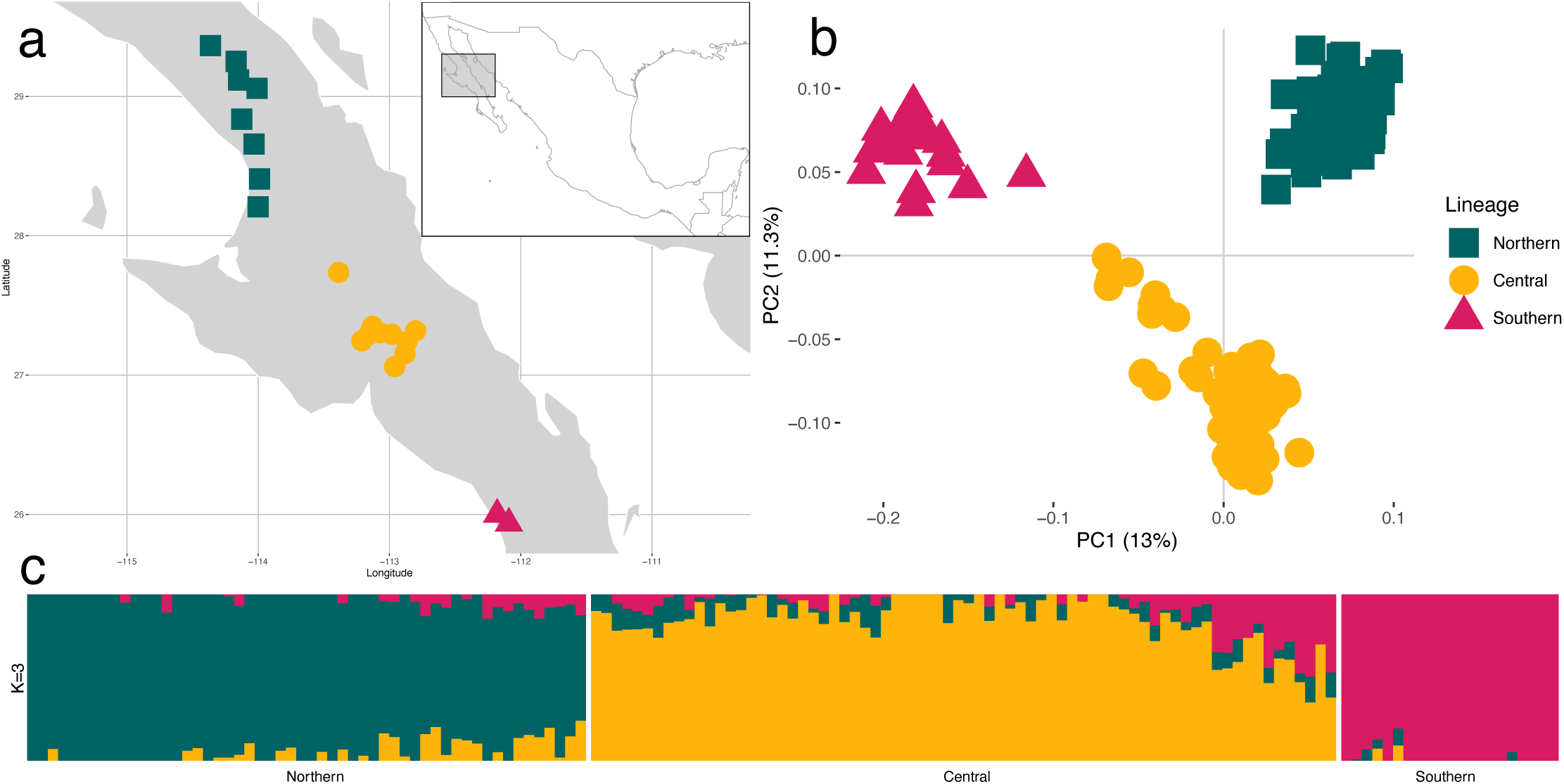
Genetic structure results of 4,411 nuclear SNPs obtained from 147 *Y. valida* samples. (a) Sampling locations (20) for this study. Shapes represent sampling locations, and colours represent genetic lineages. (b) Principal Component Analysis for PC1 and PC2. Shapes represent individuals, and colours represent genetic lineages, (c) ADMIXTURE (*K* = 3). Vertical bars represent individuals, and the genetic components are coded with different colours. Colour and shape code as the box in b: Northern (green, squares), Central (yellow, circles), Southern (pink, triangles).

Genomic DNA was converted into nextRAD sequencing libraries (SNPsaurus LLC, Oregon, USA) as in Russello et al. (2015). Briefly, DNA was fragmented with Nextera reagent (Illumina, Inc), which also ligates short adapter sequences to the ends of the fragments. The reaction was scaled to fragment 20 ng of DNA, and 50 ng of DNA were added to compensate for DNA degradation and to increase the number of fragments. The DNA was amplified for 27 cycles at 74°C. The libraries were sequenced on a HiSeq4000 150 bp single-end line (University of Oregon, Oregon, USA). The dataset was enriched with 40 raw sequences of *Y. valida* (NCBI SRA: SRR11514779) and 40 of *Y. capensis* (NCBI SRA: SRR11514778) from Arteaga et al. (2020) that were obtained for previous population genetic analyses following the same extraction and sequencing protocols described in this study; thus, representing a consistent dataset across analyses.

### Raw data processing and SNP–calling

The quality of the 160 demultiplexed raw sequences of *Y. valida* was evaluated through FastQC 0.11.9 (Andrews, 2017) and MultiQC 1.14 (Ewels et al., 2016). Adapter removal was done with custom scripts using the *bbduk* module of BBTools 1.0 (Bushnell, 2023). Clean reads were forced to 110 bp in length. A *de novo* assembly was performed in Stacks 2.6 (Catchen et al., 2013). Parameter optimization followed Mastretta-Yanes et al. (2015) and Paris et al. (2017), and 13 samples with less than 1M reads were removed as a quality control step before the assembly. We established a minimum stack depth equal to eight (*-m* 8), a maximum of three mismatches between stacks (*-M* 3), and a maximum of four mismatches between catalogue loci (*-n* 4). We called the SNPs present across the remaining 147 samples in the module populations, with a minimum genotyping rate of 0.5 and a minor allele frequency of 0.05. Hodel et al. (2017) and Shafer et al. (2017) exposed accurate resolutions on summary statistics, phylogenetic inferences, and demographic parameters using low completion cut-offs in reduced representation sequencing data; hence, we did not apply more astringent filtering to our SNP identification. When a 110 bp locus showed more than one SNP, only one variant was chosen, following the priorities in Alexander (2020) to ensure SNPs were unlinked (Pearman et al., 2022).

### Population structure

We used nuclear SNPs to assess the most likely number of genetic clusters across all samples and the membership of each sample to these clusters using two approaches: i) ADMIXTURE 1.3.0 (Alexander & Lange, 2011) was run with *K* = 1 – 20, and the optimal number of clusters was chosen using the cross-validation procedure, and ii) a Principal Components Analysis (PCA) was produced with Plink 1.90 (Purcell et al., 2007). To avoid over- or underestimating population structure (Janes et al., 2017), we verified that the assignment of samples to genetic clusters (corresponding to three lineages, see *Results*) was consistent for both approaches.

### Genetic diversity and divergence

Once the samples were allocated into discrete genetic clusters (corresponding to three lineages, see *Results*), the genetic diversity descriptors (H_E_, H_O,_ and F_IS_) for each lineage and the relative genetic divergence (paired F_ST_) between lineages were calculated with GENODIVE 3.0 (Meirmans, 2020), using the SNPs data. Data conversion was completed using PGDSpider 2.1.1.5 (Lischer and Excoffier, 2012).

To examine the isolation by distance null hypothesis, i.e., the correlation between the geographic and genetic distance between samples (Meirmans, 2012), a Mantel test was carried out using the individuals’ Euclidean genetic distance and the pairwise distance between sampling locations in meters, introducing a random jitter (±0.05°) to samples from a shared location. To evaluate the significance of the Mantel test, 99,999 permutations were used. However, as the correlation and slope values were found negligible (r = 0.25, y = 0.0001x + 32), hence rejecting the isolation by distance hypothesis, we do not present further results from this test.

### Role of selection vs drift in genetic divergence

To evaluate if population structure is a consequence of divergent selection or genetic drift, we classified the levels of genetic diversity and differentiation within and between lineages according to the evolutionary forces shaping their divergence. All the nucleotides (variant and invariant) from the sequences (110 bp long) containing the 4,411 SNPs (above) used to assess population structure were called in Stacks. The mean values of relative genetic divergence (F_ST_; Weir and Cockerham, 1984), absolute genetic divergence (d_XY_; Nei and Miller, 1990), and nucleotide diversity (π; Nei and Li, 1979) for every sequence were computed in population pairs using pixy 1.2.7 (Korunes and Samuk, 2021). Then, the correlations between pairs of summary statistics were compared. Following Shang et al. (2023), we categorized each correlation according to the relation among paired F_ST_, d_XY_, and π, given four models of adaptive divergence as expected on a genome-wide basis: 1) a positive correlation between F_ST_ and d_XY_ and a negative correlation between π and both F_ST_ and d_XY_ (divergence with gene flow); 2) uniform levels of d_XY_ across different values of F_ST_ and π, with a negative correlation between F_ST_ and π (selection under allopatry); and two models that produce a negative correlation between F_ST_ to both d_XY_ and π and a positive correlation between F_ST_ and π, characterized by 3) overall high F_ST_ with low π and d_XY_ (background selection) or 4) overall low F_ST_ with high π and d_XY_ (balancing selection). Additionally, following Kessler et al. (2023), each sequence was classified given the same four models of adaptive divergence by detecting outliers for F_ST_, d_XY_, and π between lineages (higher or lower than the 95% quantile) plus intermediate values of d_XY_ (within the 45% and 55% quantiles), and sequences were assigned to one of the models described (Appendix S1, Table S2). As neutrality was our null hypothesis, any sequence that could not be classified according to these models of divergence was identified as consistent with genetic drift.

### Outlier SNPs and SNPs associated with environmental variables

We tested the role of current ecological adaptation in promoting genetic differentiation. First, we scanned for outlier SNPs using PCAdapt 4.0.1 (Luu et al., 2017), with *K* = 3 and α = 0.1 for the false discovery rate boundary. Then, we used the Latent Factor Mixed-effect Model (LFMM, Frichot et al., 2013) to characterize the presence of SNPs with a strong association with environmental variables. LFMM was run ten times (*K* = 3) on the R package LEA 3.2.0 (Frichot and François, 2015). A total of 20 environmental variables obtained from WorldClim v1.4 (Hijmans et al., 2005) were tested for each sampling location. The Gibbs sampler algorithm was used for 10,000 cycles (burn-in = 5,000), and the median of the correlation scores (z-score) was calculated across all runs, recalibrating the frequency distribution of the p-values with lambda. We avoided any complementary correction procedures for the false discovery rate in both PCAdapt and LFMM to increase the probability of detecting outliers and SNPs correlated with environmental variables. The SNPs detected by any procedure were removed from the dataset to produce a neutral catalogue for the demographic history analyses.

### Demographic history

To understand the succession of events that could generate the observed nuclear genetic lineages, the demographic history of *Y. valida* was reconstructed using the diffusion approximation method δaδi (Gutenkunst et al., 2009). Simulations included four models of three populations developed by Portik et al. (2017): i) all three populations diverging simultaneously without any gene flow between them; one pair of populations diverging after the other from ii) North to South and iii) South to North, also without any gene flow; and iv) Northern and Southern populations diverging first, and getting into secondary contact to generate an admixed Central population. We did not include any changes in population sizes. The site frequency spectrum was computed as folded, and the best model was selected using the maximum likelihood and the Akaike information criterion (AIC).

### Putative whole chloroplast genome sequences recovery

We implemented a reference-guided workflow to reconstruct whole-chloroplast-genome sequences as per Aleman et al. (2024). The 200 raw sequences of *Y. capensis* and *Y. valida* were cleaned as above, without forcing a maximum length and establishing a minimum length = 100 bp. The clean reads were mapped to the plastome of *Y. schidigera* (GenBank: NC_032714.1) in bowtie 2.2.6 under the --*very-sensitive-local* option (Langmead and Salzberg, 2012). Mapping statistics were evaluated with the flagstat and coverage modules of SAMtools 1.15.1 (Li et al., 2009). Nucleotide calling was performed individually for each sample in ANGSD (Korneliussen et al., 2014), producing linear sequences by requiring minimum mapping and base qualities of 20, and outputting missing data as Ns *(-dofasta 2 -minMapQ 20 -minQ 20 -doCounts 1*).

### Phylogenetic relationships and divergence time

We reconstructed the intraspecific phylogenetic relationships of the samples in this study to understand their evolutionary relationships. For this, we chose the 45 Northern, 66 Central, and 19 Southern plastome sequences with higher data completeness (130 samples, maximum missing data < 0.5) to run RAxML 8.2.12 (Stamatakis, 2014), establishing *Y. schidigera* as the outgroup. A GTR + G + I model was used, letting RaxML halt automatic bootstrapping.

As the intraspecific phylogenetic analysis revealed a lack of differentiation among the plastomes of *Y. valida* (see *Results*), the divergence time of *Y. valida* and *Y. capensis* was estimated as an approximation to the age of *Y. valida*. The phylogenetic relationships of the Agavoideae subfamily were reconstructed based on the molecular clock, replicating Smith et al. (2021) in BEAST 1.8.4 (Drummond and Bouckaert, 2015). Based on Scheunert et al. (2020) and Roy et al. (2020), we produced consensus sequences for each *Y. valida* and *Y. capensis*, using all the sequences available on each species. The consensus sequences were aligned to the plastomes of 15 Agavoideae species (Appendix S1, Table S3) with MAFFT 7 default settings (McKain et al., 2016; Katoh et al., 2019). A GTR + I model was employed for the molecular clock analysis. Gamma-distributed priors for the five substitution types and uniform priors (0-1) for the base frequencies and the proportion of invariant sites were established. An uncorrelated relaxed clock model with log-normal distributed rates was set up (mean = 1, standard deviation = 0.33), assuming a Yule process for the tree prior, starting from a random tree, establishing *Hosta ventricosa* as the outgroup, and forcing the ingroup’s monophyly (Drummond and Bouckaert, 2015). A log-normal prior distribution on the age of *Yucca + Agave* was set to 14.5 million years (offset = 14.5, mean = 1, standard deviation = 1.4) based on the minimum age of the fossil *Protoyucca shadesii* (Tidwell and Parker, 1990; Iles et al., 2015). We completed two independent runs of 60 million steps with a sample every 6,000,000 steps, and the runs were joined in LogCombiner, considering a 10% burn-in for each input. A maximum credibility tree was produced with TreeAnnotator.

## RESULTS

### Raw data processing and SNP calling

A total of 400 M reads of 150 bp from 160 samples were processed (∼2.5 M reads per sample). After quality trimming and adapter removal, ∼300 M reads of 110 bp distributed in 147 samples were kept (∼2 M reads per sample). The *de novo* assembly in Stacks recovered 10,511 SNPs in 4,411 loci (minimum depth = 15x, mean missing data = 0.27, minimum sequencing quality = 39).

### Population structure, genetic diversity and divergence

Using 4,411 unlinked SNPs, the nuclear genetic structure analyses established the most likely number of lineages as three (*K* = 3) (Figure 1). The lineages were named after their geographic location as Northern, Central, and Southern, and had 54, 72, and 21 individuals, respectively. ADMIXTURE exposed evidence of substantial admixture between the Central and Southern lineages for a few Central individuals. The PCA showed some overlap along PC1 between the Northern and Central lineages, as well as the PC2 between the Northern and Southern lineages. No overlap on either axis of the PCA was seen between the Central and Southern Lineages. The overall observed (H_O_) and expected (H_E_) values of heterozygosity were 0.26 and 0.30, respectively, with a mean coefficient of inbreeding (F_IS_) of 0.17 (Appendix S1, Table S4). The relative divergence (F_ST_) between lineages was low, with the values between the Northern-Southern and Central-Southern lineages being the greatest (North-South: 0.11; North-Central: 0.05; Central-South: 0.09).

### Role of selection vs drift in genetic divergence

The patterns of correlation between measures of genetic diversity and divergence were similar for all lineage pairs, and were inconsistent with the genome-wide expectations for the four models of adaptive divergence (Figure 2): low d_XY_ with an insignificant positive correlation to F_ST_, low π with an insignificant negative correlation to F_ST_, and a significant positive correlation between d_XY_ and π (p-value < 0.05) were seen. A total of 41 sequences were consistent with one of the four models. Of those, 3 were consistent with background selection and 38 with balancing selection. A total of 4,370 sequences were identified as consistent with genetic drift.

**Figure 2.**
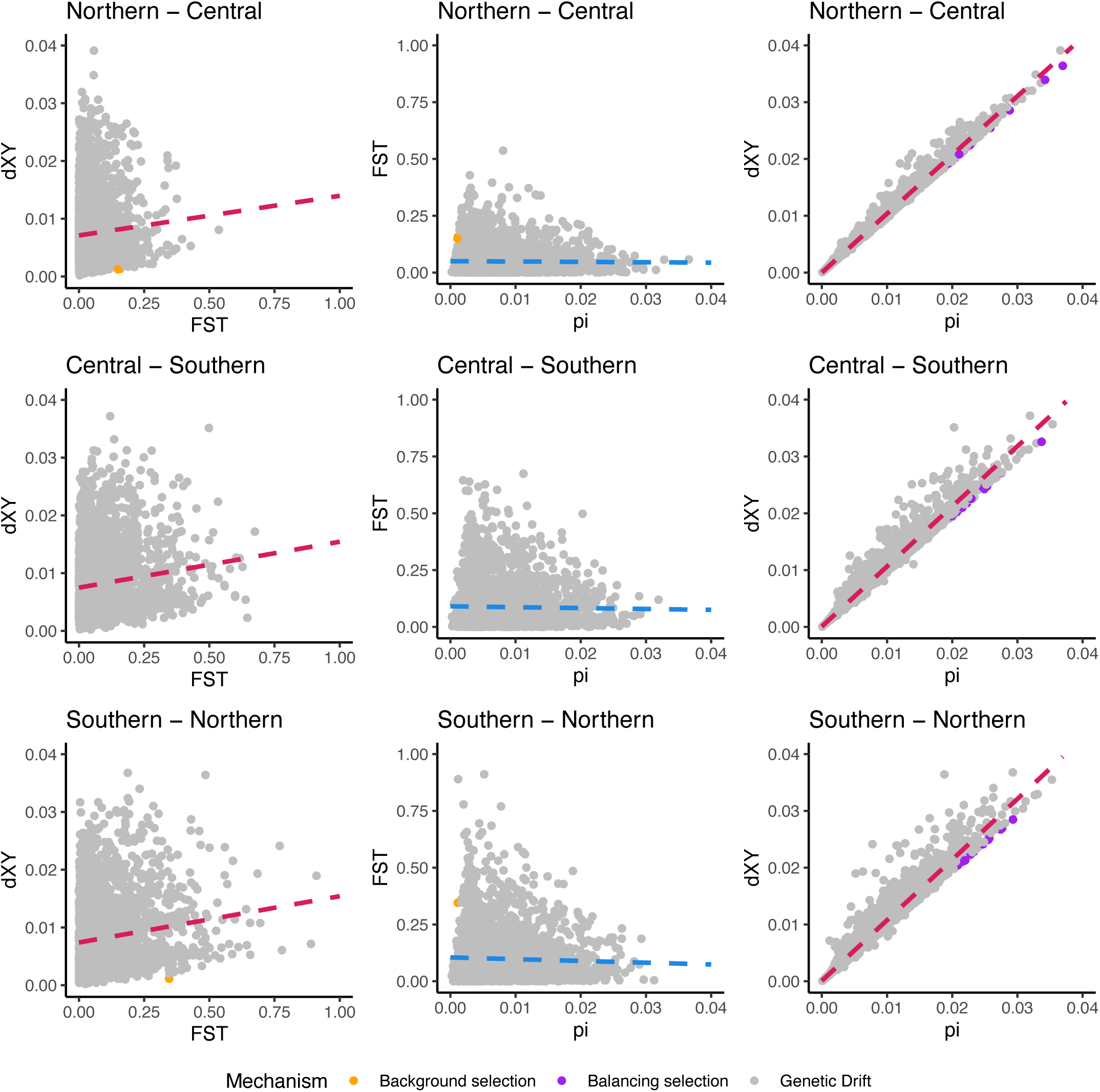
Correlation analysis for F_ST_, d_XY_ and π across 4,411 sequences (110 bp each) between pairs of nuclear genetic lineages of *Y. valida* found in this study. Each point represents a sequence, and it is coloured according to the potential evolutionary mechanism driving genetic differentiation according to the labels at the bottom. Trend lines are coloured as positive (red) or negative (blue).

### Outlier SNPs and SNPs associated with environmental variables

PCAdapt detected 20 outlier SNPs (Appendix S1, Table S5). LFMM found 63 SNPs associated with adaptation to multiple environmental variables (Appendix S1, Table S6). None of the SNPs detected by PCAdapt were also detected by LFMM, and the 83 SNPs detected were removed to produce the neutral dataset.

### Demographic history

The maximum likelihood values and the AIC indicated that the optimal demographic model was the simultaneous divergence of the three nuclear genetic lineages from a common ancestral population (Appendix S1, Table S7).

### Phylogenetic relationships and divergence time

The 130 chloroplast sequences of *Y. valida* chosen for the phylogenetic reconstruction had 4,938 segregating sites (214 without missing data) and π = 0.00096. The intraspecific phylogenetic analysis grouped the *Y. valida* plastomes into a single monophyletic clade. The three lineages of *Y. valida* could not be distinguished from the chloroplast phylogeny, and low support nodes were observed, exposing an absence of differentiation across this genome. The interspecific phylogenetic relationships and age estimates for the Agavoideae were consistent with Smith et al. (2021), and all nodes were highly supported. The molecular clock calculated an age below a million years for the last common ancestor of *Y. valida* and *Y. capensis* (Figure 3; mean ≈ 819,900; median ≈ 792,600; 95% CI ≈ 33,500 – 1,650,800).

**Figure 3.**
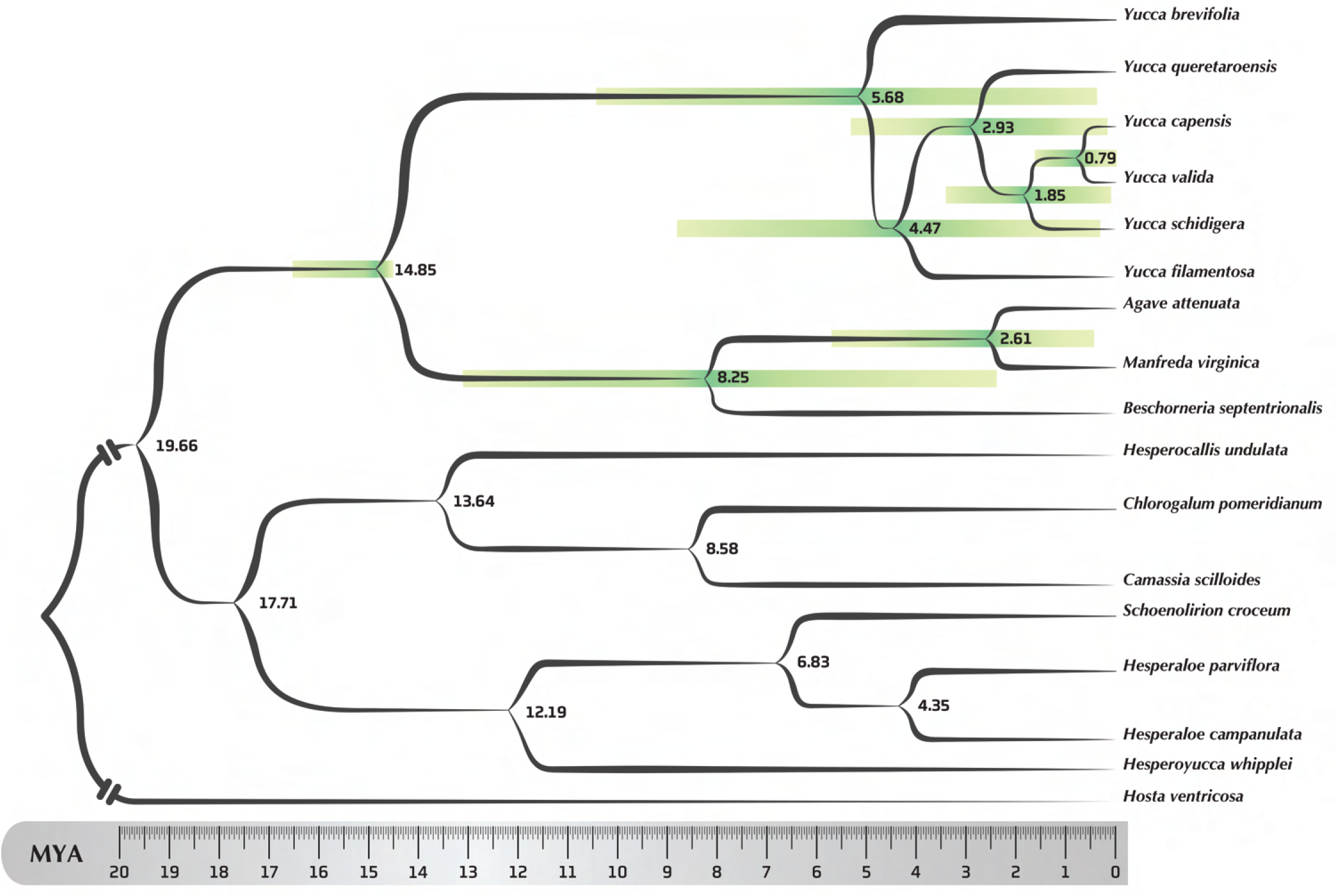
Chloroplast phylogeny of 17 Agavoideae species, estimated in BEAST 1.8.4. The contribution of this study is solely the position and divergence time of *Y. capensis* and *Y. valida*. The phylogenetic relationships among the chloroplast genome sequences within the Agavoideae are discussed in Smith et al. (2021). The tree was produced with *Hosta ventricosa* at the root and calibrated with a log-normal distribution on the age of *Yucca* and *Agave* at 14.5 million years. Numbers represent the estimated medians of the nodes’ divergence times as millions of years from now. Horizontal bars represent the 95% confidence intervals. Branch support on every node = 1.

## DISCUSSION

We examined the phylogeography of *Y. valida* throughout its range and identified three monophyletic nuclear lineages accompanied by a single chloroplast lineage. The age estimated for the last common ancestor of *Y. valida* and its sister species was below 1 MYA, suggesting that the nuclear differentiation in *Y. valida* is a consequence of recent events. While the time of origin of the three nuclear lineages is uncertain, ecological niche modelling has predicted that during the Last Interglacial, the area with suitable conditions for *Y. valida* was completely absent (Arteaga et al., 2020). Climatic oscillations are not the only possible explanation for the observed patterns of divergence (Dolby et al., 2022). Low dispersal without a climatic barrier can also result in genetic differentiation (Araya-Donoso et al., 2022); thus, the role of restricted gene flow should be considered. Likewise, the absence of an isolation by distance pattern indicates that divergence cannot be attributed solely to the distance between populations. Collectively, this suggests that habitat fragmentation caused by climatic changes, low dispersal, and an extensive geographical range gap have acted as cumulative mechanisms, leading to allopatric divergence, as indicated by the patterns of genetic differentiation consistent with drift.

The values of genetic differentiation between the three nuclear lineages were low (mean paired F_ST_ = 0.08). Estimates of genetic differentiation between lineages (mean F_ST_ = 0.08) agreed with estimates based on microsatellite loci for *Y. valida* (F_ST_ = 0.059; Alamo-Herrera et al., 2022), *Y. schidigera* (F_ST_ = 0.067; De la Rosa-Conroy et al., 2020), and *Y. brevifolia* (F_ST_ = 0.061; Starr et al., 2013), and are greater than those in its sister species, *Y. capensis* (F_ST_ = 0.022; Luna-Ortiz et al., 2022). The low levels of differentiation can result from potentially large historical effective population sizes (Arteaga et al., 2020) and reflect some level of gene flow between lineages. Estimates of the dispersal rates indicate that pollen movement in *Y. valida* is approximately 50 m (Alamo-Herrera et al., 2022), and seed-mediated dispersal also occurs at short distances (Waitman et al., 2012), which could maintain connectivity between populations through a stepping-stone model. In this sense, the presence of genetic structure, the patterns of genetic diversity and divergence, and the low differentiation coincide with contemporary genetic drift to be the driving force of divergence in this species.

Genetic diversity was similar across the three lineages. The diversity estimates (H_O_ = 0.26, H_E_ = 0.31) were higher than those described for this species with the same type of genetic markers (H_O_ = 0.11, H_E_ = 0.17; (Arteaga et al., 2020)). The disparity between these values reiterates the effect of different methodological variables on SNP calling in population genomics (O’Leary et al., 2018). Arteaga et al. (2020) assembled loci from samples of two species —*Y. valida*, *Y. capensis*, and their hybrid. Here, we generated a SNP dataset with samples of a single species, thus modifying the number of polymorphisms detected. SNP-calling among differentiated taxa implies retrieving alleles fixed for a species or a population that are polymorphic between species or populations, affecting diversity estimates (Shafer et al., 2017); this reiterates that conclusions based on genetic diversity estimates should be made on a case-by-case basis (Sopniewski and Catullo, 2024). The inbreeding coefficient (F_IS_ = 0.17) was also lower than in Arteaga et al. (2020) (F_IS_ = 0.35). The high diversity and low inbreeding detected in this work are congruent with hypothetical high historical population sizes and with the reproductive system of *Yucca*, which favours outcrossing and limits self-pollination (Massey and Hamrick, 1999; Huth and Pellmyr, 2000; Marr et al., 2000; Alamo-Herrera et al., 2022).

The demographic modelling using nuclear data supported the hypothesis of a simultaneous divergence among the three lineages without gene flow. The lack of intraspecific differentiation in the plastome of *Y. valida* indicates that the drivers of genetic divergence in *Y. valida* have been accumulating long enough to be recorded in the nucleus and so recently not to be registered in the chloroplast. The lack of congruence between outlier SNPs and SNPs associated with environmental variables, plus the absence of signals of adaptive divergence, rule out the role of selection as the driver of genetic divergence among the three lineages of *Y. valida*; however, this does not mean that the species is not adapted in any way to its local environment. Our results support the role of drift under allopatry as the driver of genetic divergence of plant species across the BCP.

## CONCLUSIONS

Determining the impact of the drivers of species’ genetic diversity and differentiation is a crucial aspect of evolutionary biology. Here, we tested the role of drift vs selection in promoting divergence on a representative taxon of the BCP. The history of *Y. valida* is one of allopatric differentiation. Addressing the origin and colonization of *Yucca* in the Peninsula remains an essential research gap in the evolutionary history of this genus.

## ACKNOWLEDGEMENTS

We respectfully acknowledge that the territory where the samples were collected are the ancestral homelands of the Cochimí people. This study was completed at Centro de Investigación Científica y Educación Superior de Ensenada, which is on the traditional territory of the Ku’ahles, Cochimíes, Pa ipais and Kiliwas, to whom we show our respect. The authors thank Cynthia Rocío Álamo-Herrera, Leonardo de la Rosa-Conroy, Astrid Luna-Ortiz, Mario Salazar, José Delgadillo, and Lita Castañeda for their contributions in the laboratory and the field. This work was supported by Consejo Nacional de Humanidades, Ciencias y Tecnologías (CB-2014-01-238843, infra-2014-1-226339), from which Alberto Aleman also received a master’s degree scholarship (Grant 748805). The Rufford Foundation (RSG 13704-1) and the JiJi Foundation provided financial support for part of this study. We are also grateful to the Associate Editor and the anonymous reviewers for their valuable comments and suggestions, which helped improve our manuscript’s quality. Finally, we thank Camille Kessler and Flavia Termignoni-García for their bioinformatic advice and Enrique Ruiz for his work in Fig. 3.

## AUTHOR CONTRIBUTIONS

Conceptualisation: MCA. Developing methods, conducting the research, data interpretation and writing: AA, JGP, MCA, and RBB. Data analysis and preparation of figures and tables: AA. All authors contributed to the manuscript and approved its final version.

## DATA AVAILABILITY

Data and scripts for this study can be found at https://github.com/al-aleman/datillilo. High-throughput sequencing raw data is available at https://datadryad.org/stash/share/Ujo8Mi1d2g50_VAAsdXzMWKgsYcffCGD59_VLFGyc3U and https://doi.org/10.5061/dryad.jh9w0vtgv.

## SUPPORTING INFORMATION

Additional Supporting Information may be found online in the supporting information section at the end of the article:

**APPENDIX S1.** Supplemental tables.

## Notes

### Competing Interest Statement

The authors have declared no competing interest.

### Summary of Updates

In addition to the modifications prompted by reviewers, we have made some adjustments in terms of the paper's wording and structure, which is aimed at further enhancing the readability and flow of the document. These changes solely pertain to the paper's form and do not impact its content or the core findings.

