## Appendix S1 for "Divergent lineages in a young species: the case of Datilillo (*Yucca valida*), a broadly distributed plant from the Baja California Peninsula"

**Table S1.** Site IDs, approximate coordinates, and number of samples (n) collected across the distribution of *Y. valida* for this study. The number of samples includes 40 individuals collected by Arteaga et al. (2020) used to enrich the final dataset.

| Site ID | Latitude | Longitude | n |
| --- | --- | --- | --- |
| 100 | 29.36467 | -114.36412 | 3 |
| 16 | 29.25202 | -114.16822 | 6 |
| 15 | 29.11758 | -114.14969 | 6 |
| 26 | 29.05869 | -114.00852 | 5 |
| 105 | 28.83702 | -114.12524 | 10 |
| 104 | 28.65867 | -114.03167 | 4 |
| 103 | 28.40771 | -113.99003 | 9 |
| 14 | 28.20819 | -114.00066 | 11 |
| 1 | 27.73669 | -113.38708 | 12 |
| 13 | 27.31694 | -112.79994 | 4 |
| 7 | 27.31363 | -113.12936 | 8 |
| 8 | 27.30480 | -113.07413 | 5 |
| 9 | 27.2925 | -112.98147 | 5 |
| 4 | 27.29052 | -113.15716 | 11 |
| 6 | 27.24591 | -113.21036 | 7 |
| 10 | 27.23780 | -112.86741 | 2 |
| 102 | 27.15399 | -112.88222 | 6 |
| 101 | 27.06119 | -112.96024 | 12 |
| 90 | 26.000366 | -112.17968 | 6 |
| 91 | 25.936442 | -112.09022 | 15 |

**Table S2.** Four models of adaptive divergence used to test the role of selection vs drift in genetic divergence. Each of the 4,411 sequences of 110 bp was classified according to the following thresholds for F_ST_, π and d_XY_.

| Model | F_ST_ | π | d_XY_ |
| --- | --- | --- | --- |
| Divergence with gene flow | > 0.95 | < 0.05 | > 0.95 |
| Selection under allopatry | > 0.95 | < 0.05 | > 0.45 and < 0.55 |
| Background selection | > 0.95 | < 0.05 | < 0.05 |
| Balancing selection | < 0.05 | > 0.95 | > 0.95 |

**Table S3**. Taxa and GenBank accessions used to investigate the phylogenetic relationships and time of origin of *Y. valida* for this study.

| Taxon | Accession No. |
| --- | --- |
| *Agave attenuata* | NC_032696.1 |
| *Beschorneria septentrionalis* | NC_032699.1 |
| *Camassia scilloides* | NC_032700.1 |
| *Chlorogalum pomeridianum* | NC_032701.1 |
| *Hesperaloe campanulata* | NC_032702.1 |
| *Hesperaloe parviflora* | NC_032703.1 |
| *Hesperocallis undulata* | NC_032704.1 |
| *Hesperoyucca whipplei* | NC_032705.1 |
| *Hosta ventricosa* | NC_032706.1 |
| *Manfreda virginica* | NC_032707.1 |
| *Schoenolirion croceum* | NC_032710.1 |
| *Yucca brevifolia* | NC_032711.1 |
| *Yucca filamentosa* | NC_032712.1 |
| *Yucca queretaroensis* | NC_032713.1 |
| *Yucca schidigera* | NC_032714.1 |

**Table S4.** Global genetic diversity of *Y. valida* and by lineage in 4,411 nuclear SNPs. Number of individuals per lineage (n) and genetic diversity descriptors (H_O_, observed heterozygosity; H_E_ expected heterozygosity; F_IS_, inbreeding coefficient).

| Lineage | n | *H_O_* | *H_E_* | *F_IS_* |
| --- | --- | --- | --- | --- |
| Northern | 54 | 0.25 | 0.30 | 0.17 |
| Central | 72 | 0.26 | 0.30 | 0.17 |
| Southern | 21 | 0.26 | 0.32 | 0.17 |
| Global | 147 | 0.26 | 0.31 | 0.17 |

**Table S5.** Outliers detected by PCAdapt from 4,411 nuclear SNPs. CHR and POS refer to the SNP coordinates given by Stacks during the *de novo* assembly.

| CHR | POS |
| --- | --- |
| 2717 | 43 |
| 4154 | 87 |
| 6056 | 7 |
| 6773 | 17 |
| 10893 | 51 |
| 11117 | 70 |
| 13210 | 79 |
| 15347 | 6 |
| 17144 | 9 |
| 17635 | 39 |
| 18085 | 46 |
| 19935 | 54 |
| 21656 | 70 |
| 25913 | 27 |
| 450329 | 53 |
| 600342 | 56 |
| 611441 | 60 |
| 630707 | 101 |
| 1074414 | 80 |
| 1379936 | 80 |

**Table S6.** Sites associated with adaptation to environmental variables detected by LFMM from 4,411 nuclear SNPs. CHR and POS refer to the SNP coordinates given by Stacks during the *de novo* assembly.

| CHR | POS |
| --- | --- |
| 241 | 40 |
| 769 | 47 |
| 2825 | 11 |
| 3323 | 58 |
| 3427 | 13 |
| 3538 | 86 |
| 4069 | 91 |
| 4158 | 86 |
| 6130 | 40 |
| 7398 | 72 |
| 7458 | 84 |
| 7931 | 80 |
| 8838 | 38 |
| 9691 | 62 |
| 11751 | 11 |
| 13524 | 97 |
| 13935 | 78 |
| 14780 | 58 |
| 16872 | 94 |
| 17498 | 66 |
| 17860 | 100 |
| 18484 | 49 |
| 19262 | 48 |
| 20266 | 56 |
| 20826 | 40 |
| 20960 | 57 |
| 21412 | 97 |
| 22952 | 31 |
| 23755 | 79 |
| 23960 | 24 |
| 25568 | 89 |
| 25687 | 42 |
| **CHR** | **POS** |
| 25693 | 50 |
| 27349 | 102 |
| 28414 | 6 |
| 30742 | 67 |
| 32927 | 30 |
| 35014 | 33 |
| 38388 | 92 |
| 38681 | 90 |
| 40849 | 59 |
| 46914 | 42 |
| 52185 | 28 |
| 54121 | 14 |
| 69932 | 71 |
| 70103 | 101 |
| 75359 | 14 |
| 75650 | 37 |
| 132662 | 67 |
| 302866 | 95 |
| 385172 | 32 |
| 509397 | 5 |
| 510503 | 79 |
| 535444 | 50 |
| 774028 | 37 |
| 778758 | 20 |
| 803977 | 85 |
| 1153982 | 13 |
| 1162120 | 19 |
| 1226455 | 75 |
| 1413338 | 11 |
| 1429185 | 27 |
| 1509272 | 41 |

**Table S7.** Summary of the demographic models tested to understand the events that could generate the three nuclear genetic lineages of *Y. valida*, reconstructed using δaδi and Daniel Portik's δaδi_pipeline (Portik et al. 2017). The best five scoring replicates from each model are shown, ranked by the Akaike information criterion (AIC). Simulations did not include any changes in population sizes.

| Model | AIC | log-likelihood |
| --- | --- | --- |
| All three populations diverging simultaneously without any gene flow between them | 7064.64 | -3528.32 |
|  | 7070.54 | -3531.27 |
|  | 7073.34 | -3532.67 |
|  | 7078.68 | -3535.34 |
|  | 7079.66 | -3535.83 |
| Northern and Southern populations diverging first and getting into secondary contact to create the Central population | 7320.32 | -3654.16 |
|  | 7330.18 | -3659.09 |
|  | 7355.94 | -3671.97 |
|  | 7357.2 | -3672.6 |
|  | 7359.58 | -3673.79 |
| One pair of populations diverging after the other from South to North, without any gene flow | 7741.64 | -3864.82 |
|  | 7777.88 | -3882.94 |
|  | 7877.72 | -3932.86 |
|  | 8002.3 | -3995.15 |
|  | 8017.94 | -4002.97 |
| One pair of populations diverging after the other from North to South, without any gene flow | 9573.48 | -4780.74 |
|  | 9786.98 | -4887.49 |
|  | 9815.02 | -4901.51 |
|  | 9820.98 | -4904.49 |
|  | 9904.06 | -4946.03 |
